# Single rod-shaped cell fluctuations from stochastic surface/volume growth rates

**DOI:** 10.1101/852624

**Authors:** Orso Maria Romano, Marco Cosentino-Lagomarsino

## Abstract

Growing rod-shaped bacterial cells need to modulate the production rates of different surface and bulk components. Population data show that the balance between these rates is central for cell physiology, and affects cell shape, but we still know little about these processes in single cells. We study a minimal stochastic model where single cells grow by two fluctuating volume-specific surface and volume growth rates, solving for the steady-state distributions and the correlation functions of the main geometric features. Our predictions allow us to address the detectability of different scenarios for the intrinsic coupling between the allocation of resources to surface and bulk growth.

## 1 Introduction

The rigorous study of the shapes of living organisms has been crucial to the development of biology as a modern science [1, 2, 3]. In bacteria, like in higher organisms, shape-related traits play a role in mediating fitness and adaptation [4, 5]. To a first approximation, the shape of a bacterial cell can be described by its size-related features such as length and width. Today, advances in microscopy allow us to access the dynamics of these variables and to monitor cell-to-cell variability of growing bacterial cells [4, 6]. This opens the question of how growth and division dynamically cast cell shape, linking shape homeostasis and growth control to the physiology of single cells.

Two processes contribute to the determination of individual bacterial cell shape: growth and division. While the relationship between bacterial size control and the timing of cell division is being thoroughly investigated [7, 8, 9], the problem of how single microbial cells achieve shape homeostasis while growing between two successive divisions is comparatively less explored [10, 11, 12, 13], and can be – to some extent – factored out from that of regulation of cell division. In particular, the relationship between cell growth and shape control has to depend on the differential biosynthesis of cellular mass (volume) and surface-specific “building blocks” [14]. A pivotal question is how energy is allocated by the cell into bulk mass growth vs the growth of surface components. The answer to this question has to depend on several processes, including the allocation of biosynthetic resources [15], direct regulatory mechanisms sensing cell mechanics [12], cellular metabolism [16].

While we start gaining some insight in such processes, our knowledge is still too limited for a comprehensive description of their implication in shape homeostasis. A less ambitious approach is to study simplified models able to describe the relevant dynamics through a simple mathematical formalism dealing with effective variables. A remarkable example is provided by a recent study by Harris and Theriot, which proposes a “relative rates” model for the growth of rod-like bacteria [11]. This model captures the dynamics of the average cell shape by assuming that both cellular volume and surface production within one cell cycle are proportional to cell volume through constant, environment-specific rates. As further studies pointed then out, the relation between such surface/volume growth rates can be considered as proxies of how the cell machinery partitions its proteome between bulk vs surface biosynthesis [17].

Here, we analyze a stochastic extension of the Harris and Theriot model to describe shape fluctuations at the single-cell level. This model is obtained by adding coupled white-noise fluctuations to the volume- and surface-specific growth rates defined in ref. [11], and is formulated as Langevin equations describing the correlated dynamics of cell-to-cell variability in shape-related observables (e.g. width, length, and the surface-to-volume ratio). This model can make predictions on how single rod-like bacterial cells achieve their (environment-dependent) shape homeostasis through growth and on how they respond to growth perturbations (such as nutrient shifts). We find that – although the growth-rate fluctuations set a multiplicative noise term in the Langevin equations – the stationarity, the steady-state distributions, and the auto- and cross-correlation functions of the geometric observables across the population are well predicted by approximating the width fluctuation dynamics with that of an Ornstein-Uhlenbeck process. Our results furthermore show that an intrinsic coupling between the temporal fluctuations in the two growth rates would affect the cross-correlation functions of the fluctuations in geometry-related observables such as cell width and length. We interpret this coupling as related to the aforementioned allocation problem of cellular resources into bulk vs surface biosynthesis.

## 2 Model

The Harris-Theriot model [11] is based on the assumption that, in an “average” rod-shaped cell, the instantaneous rates of surface (*S*) and volume (*V*) synthesis both scale linearly with volume, i.e.,

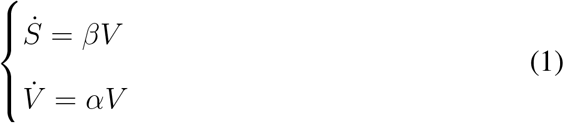

where the dots indicate time derivatives. The parameter *α* (dimensionally, an inverse time) is the specific rate of volume growth, while *β* the (volume-specific) rate of surface material synthesis (with units of inverse length times inverse time). With few, simple ingredients, this model reproduces how the average surface-to-volume ratio *S/V* (an effective proxy for shape) relaxes and settles to nutritional shifts and perturbations of cell-wall synthesis [11].

The model analyzed here extends the Harris & Theriot model to single cells. As in the original model, it describes the growth dynamics that occurs within consecutive cell divisions (factoring out the role of divisions in cell shape [8]). For simplicity, we approximate the rod-shaped cell by a cylinder, neglecting the caps (Fig. 1). This approximation is reasonable when dealing with surface and volume *variations*, since caps are synthesized at cell division and their interdivision growth is negligible. Under these assumptions volume and surface are simple functions of the two linear dimensions width *w* and length *L* (we neglect for simplicity geometric factors *π* and *π*/4, respectively):

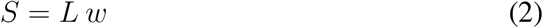

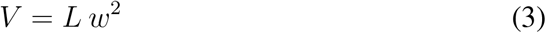

**Figure 1:**
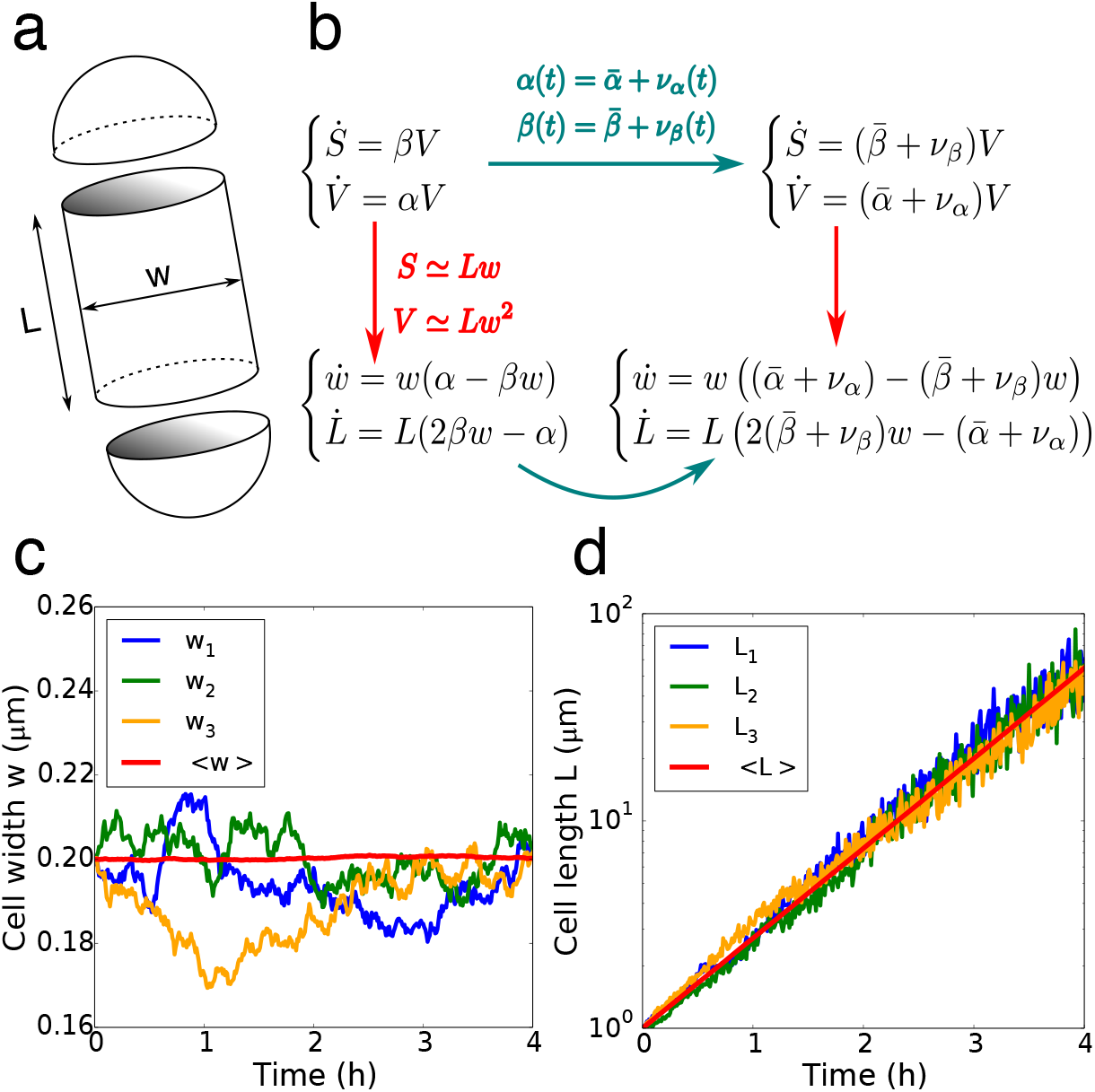
Basic definitions and simulations of the single-cell growth model for rod-shaped cells. **(a)**: Cellular surface *S* and volume *V* are represented as those of a cylinder of width *w* and length *L*, neglecting caps. **(b)**: The mathematical model introduces temporal fluctuations to the volume and surface growth rates *α, β* of the Harris-Theriot model (top left). The resulting stochastic equations are equivalently expressed by width and length (bottom right) or surface and volume (top right). **(c)**: Cellular width time series obtained by simulating the first of Eqs. 7. The red curve represents the average width time series over 2000 independent realizations. **(d)**: Cell length time series obtained by simulating the second of Eqs. 7, in logarithmic scale. The red curve represents the average length time series over 2000 independent realizations. Parameters: 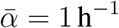, *σ_α_* = 0.05 h^−1^, 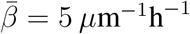, *σ_β_* = 0.25 *μ*m^−1^h^−1^, *ρ* = 0.

In other words, under the shape constraint *V* = *S w* Eqs. 1 are equivalent to the following system of ODEs:

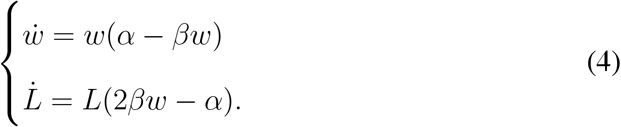

To define a single-cell growth model, we assume the rates *α* and *β* to be the result of the stochasticity of cellular processes, and thus characterized by average values 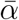 and 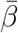 and temporal fluctuations *ν_α_* = *ν_α_*(*t*) and *ν_β_* = *ν_β_*(*t*),

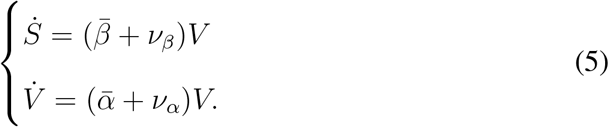

We model fluctuations *ν_α,β_* as zero-average, delta-correlated (white) noises, i.e.

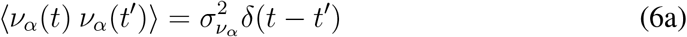

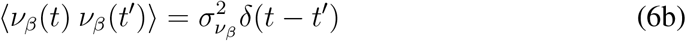

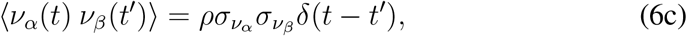

where the noise amplitudes 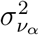 and 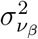 are in *T*^−1^ and *L*^−1^*T*^−1^ units, respectively, thus corresponding to the volume- and surface-specific growth rates’ empirical standard deviations normalized by the unitary time step.

The correlation coefficient *ρ* represents the assumption that the surface and volume (bulk) rates could *a priori* be coupled by cell physiology, distinguishing between scenarios in which fluctuations *ν_α_, ν_β_* are (i) completely uncorrelated (*ρ* = 0), (ii) negatively correlated (*ρ* ∈ [−1, 0)), e.g. by competition between surface and volume synthesis, or (iii) positively correlated (*ρ* ∈ (0, 1]), e.g. by common allocation of resources for bulk and surface growth. Please note that, while in the original average-cell model volume growth is typically interpreted as bulk (mass), and mass density as a constant (on average), the same interpretation in our single-cell extension corresponds to the assumption of constant density for single cells. This assumption can be relaxed by model extensions explicitly describing the dynamics of mass and volume, or of mass and surface, which we did not address in this study.

Under these assumptions, the stochastic version of Eqs. 4 write

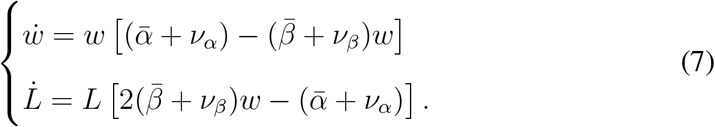

We observe that, as in the deterministic model (Eqs. 4), the equation for width is autonomous, while the dynamics of length *L* is coupled to width *w*. For steady-state values of the average width 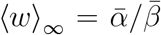, mean length increases exponentially with rate equal to the volume growth rate, 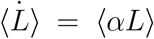. Neither *L* nor *q* = log(*L*/*L*_0_) converge to a finite average value in the limit of large times, meaning that their fluctuations *δL* and *δq* are not stationary – which suggests 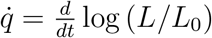 as a more appropriate observable, which might reach stationarity.

## 3 Results

We are interested in understanding how, in the stochastic model, the volume- and surface-growth rate fluctuations *ν_α_, ν_β_* and their istantaneous correlation *ρ* affect cell-to-cell variability in cell width *w*, logarithmic length growth 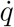 and surface-to-volume ratio *z* = *S/V*. We chose to focus our attention on such (stationary) variables as they are those that are typically probed in single-cell experiments, for example by the use of microfluidic devices. Our goal is to characterize the steadiness, fluctuation behavior, and auto- or cross-correlation functions of such geometric observables.

### The Langevin equations for the temporal dynamics of fluctuations of shape-related variables include both additive and multi-plicative noise terms

Width fluctuations can be expressed as the deviation *δw*(*t*) = *w*(*t*) – 〈*w*〉_∞_, around the steady-state mean value. For small fluctuations, the expression of the temporal dynamics of *δw* can be obtained from the linearization of the first of Eqs.4:

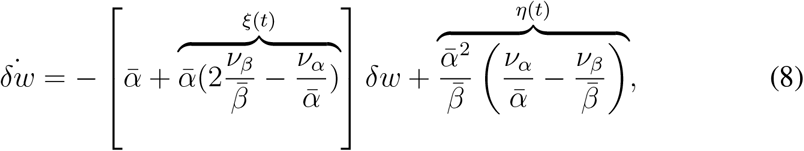

which comprises one additive and one multiplicative source of noise, here called *η*(*t*) and *ξ*(*t*), respectively.

Equation 8 can be seen as the equation for the dynamics for an overdamped point particle subject to thermal noise (the additive noise) in a harmonic potential whose stiffness is fluctuating (multiplicative noise),

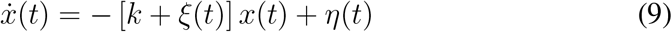

whose solution is not trivial. When *ξ*(*t*) = 0 ∀t, Eq. 9 becomes

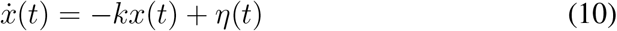

which is the Langevin equation for an Ornstein-Uhlenbeck process (overdamped Brownian particle in a harmonic potential) [18]. When *η*(*t*) = 0 ∀*t*, Eq.9 becomes

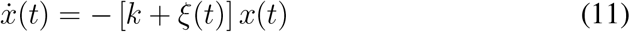

which is the Langevin equation for a multiplicative random walk with drift (Black-Scholes model) [18].

Both additive and multiplicative noises in Eq. 8 can be expressed in terms of the mean values of the volume and surface growth rates 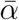, 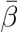 and of their relative fluctuations 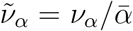, 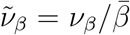 (corresponding to the actual sources of noise of the model):

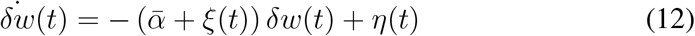

with

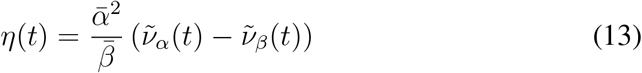

and

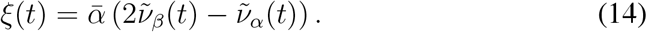

Both *η* and *ξ* are zero-average. Their variances depend on the correlation coefficient *ρ*:

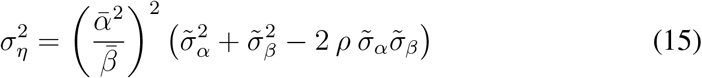

and

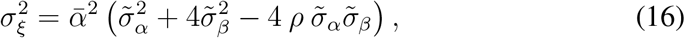

where 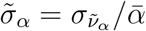 and 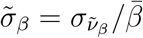.

The SDE describing the fluctuations in the surface-to-volume ratio *δz*(*t*) = *z*(*t*) – 〈*z*〉_∞_, with *z* = *S/V* = 1/*w*, can be obtained exactly since the equation ruling the dynamics is linear:

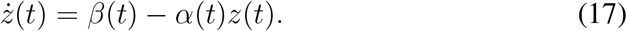

The surface-to-volume ratio fluctuations obey the following Langevin equation, again characterized by both additive and multiplicative noise terms:

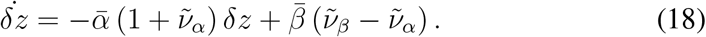

Finally, to evaluate length fluctuations we combine Eqs. 4 with Eq. 8, obtaining

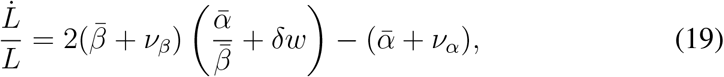

which can be written as

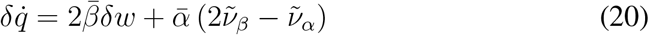

where *q* = log(*L/L*_0_), hence 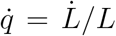 and 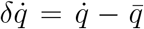 with 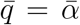. Equation 20 shows that the fluctuations around exponential length growth have no restore force, are coupled to width, and are subject to additive noise.

### Higher multiplicative noise produces wider and more skewed steady-state distributions of width fluctuations (Fig. 2)

**Figure 2:**
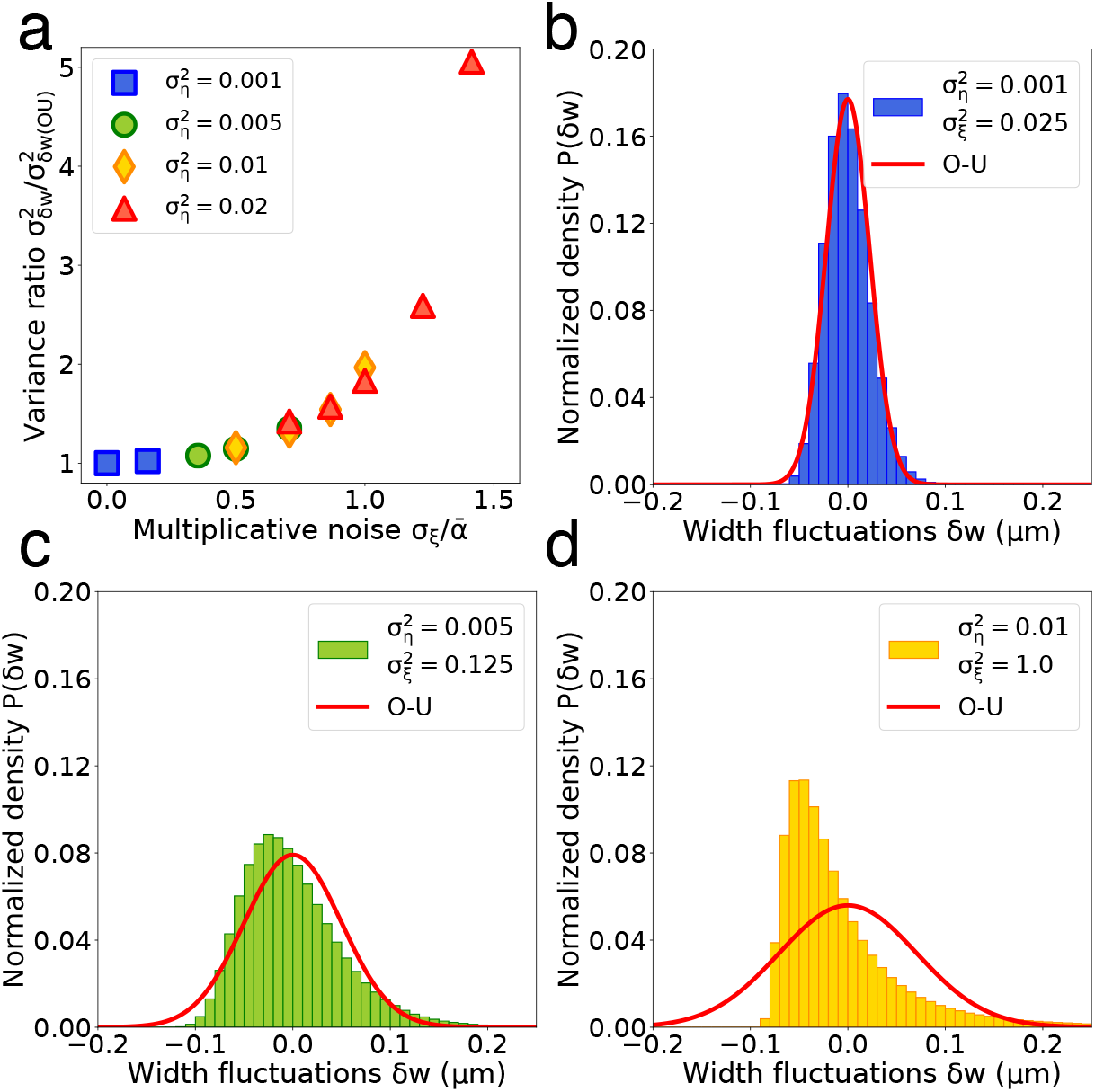
Higher multiplicative noise produces wider and more skewed steady-state distributions of width fluctuations. **(a)**: The ratio of the variance of width fluctuations at steady state and the prediction of the Ornstein-Uhlenbeck approximation (negligible multiplicative noise regime) increases superlinearly with *σ_ξ_* at fixed 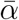. Panels b-d correspond to additive noise levels with increasing variances 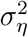, resulting in wider and more skewed distributions with respect to the pure Ornstein-Uhlenbeck regime. **(b-d)**: Comparison of steady-state width distributions with the Ornstein-Uhlenbeck prediction (red line) in the low (≃ 16% of the average volume growth rate 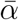, panel b), intermediate (≃ 35% of 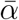, panel c) and high multiplicative noise (=100% of 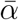) regimes. Parameters – **(b)**: 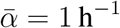, *σ_α_* = 0.16 h^−1^, 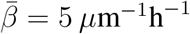, *σ_β_* = 0 *μ*m^−1^h^−1^; **(c)**: 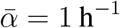, *σ_α_* = 0.35h^−1^, 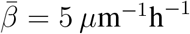, *σ_β_* = 0 *μ*m^−1^h^−1^; **(d)**: 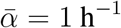, *σ_α_* = 0 h^−1^, 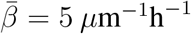, *σ_β_* = 2.5 *μ*m^−1^h^−1^.

For small multiplicative noise (measured by 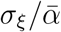, where the amplitude of the noise *ξ* can be obtained from Eq. 16 in the case of width fluctuations), the Langevin equation for the width fluctuation dynamics (Eq. 8) is well approximated by that of an Ornstein-Uhlenbeck process. In this regime, width fluctuations reach stationarity on a time scale set by 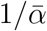. The steady-state distribution of width fluctuations is a zero-averaged Gaussian distribution whose variance is given by

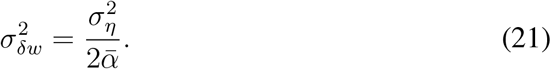

The Ornstein-Uhlenbeck approximation remains valid as long as 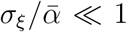. Values of *σ_ξ_* comparable to, or higher than 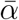 produce increasingly wider and more skewed distributions with respect to the variance set by the additive noise only (Fig. 2). In particular, relative deviations from such Ornstein-Uhlenbeck prediction follow a superlinear law with 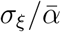 (Fig. 2a).

It should be noted that when *ρ* = 0, given that variations in the additive and multiplicative noise terms in the Langevin equations for width fluctuations can only be obtained through a change in the same four parameters 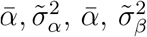, only a fraction of all the possible variances of *ξ* are accessible for a fixed variance of *η*, being the variances of *η* and *ξ* linked by the constraint

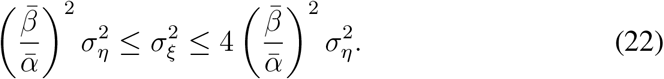

### Increasingly positive correlations between surface and volume growth-rate fluctuations dampen width fluctuations at steady state, while increasingly negative ones increase them

Equations 15 and 16 express the variances of additive and multiplicative noise terms for the Langevin equation of the dynamics of width fluctuations in terms of the same five parameters: the average volume- and surface-growth rates 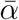, 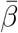, their coefficients of variation 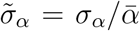, 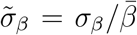, and the correlation coefficient of volume- and surface-growth rates fluctuations as defined in Equation 6c. Both 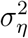 and 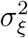 linearly decrease with *ρ*.

Since the levels of steady-state width fluctuations 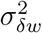 in the low multiplicative noise regime are proportional to the additive noise variance (Eq. 21), Equation 15 also implies that the more correlated are the fluctuations of *α*(*t*) and *β*(*t*), the less disperse are the width fluctuations at steady state (Fig. 3). In particular, for *ρ* = 1 (i.e. when the two volume- and surface-specific growth rates are subject to the same white noise) the variance of *δw* at steady state is at its minimum, its maximum being attained for complete anticorrelation *ρ* = –1. If the fluctuations of *α*(*t*) and *β*(*t*) are characterized by the same relative standard deviations 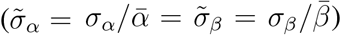, the minimum is zero (cf. Eq. 15). The slope of the linear decrease of 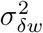 with *ρ* is function of both mean and coefficient of variation of the two growth rates 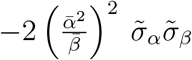 (Fig. 3a).

**Figure 3:**
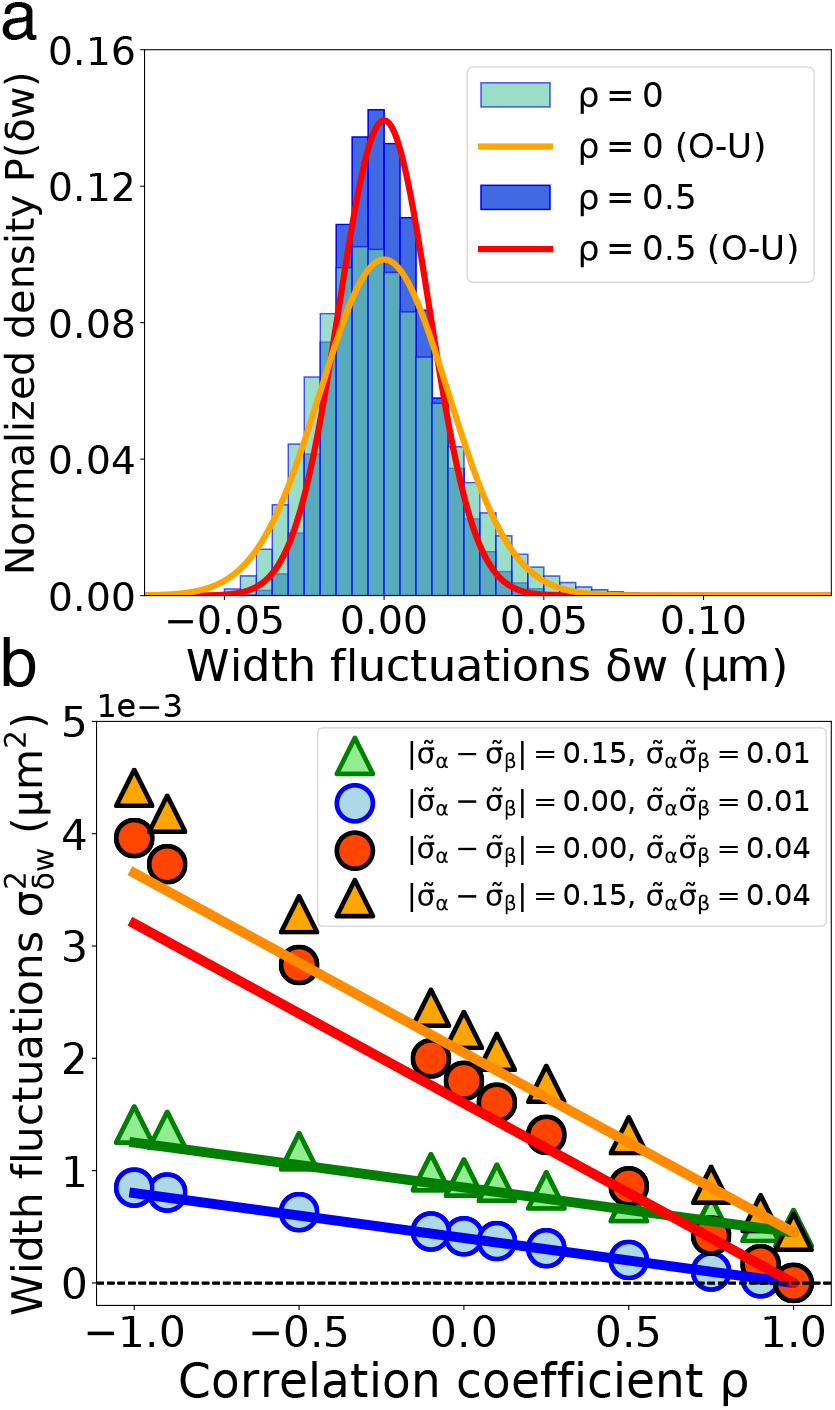
Increasing correlation between surface- and volume-growth rate fluctuations decreases the variance of width fluctuations at steady state. **(a)**: Direct comparison between steady-state distributions of width fluctuations at different values of the correlation coefficient between volume- and surface growth rates *ρ*, all other parameters equal (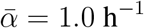, *σ_α_* = 0.1h^−1^, 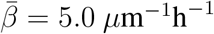, *σ_β_* = 0.5 *μ*m^−1^h^−1^). The red and orange lines indicate the Ornstein-Uhlenbeck theoretical prediction (which neglects correlations but is *ρ*-dependent through the noise amplitudes), given Eqs. 21 and 15) for *ρ* = 0 and *ρ* = 0.5, respectively. Both predictions are good, being in the low multiplicative noise regime (5% and 3% of 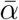, respectively). **(b)**: Variance of width fluctuations at steady state as a function of *ρ*, correlation coefficient of the volume- and surface growth rates, for different combinations of the amplitudes of volume- and surface-specific growth rates fluctuations 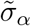, 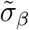. The difference between the two 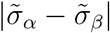 sets the minimum 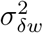 at steady state, attained for *ρ* = 1. Indeed, for 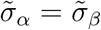, width fluctuations vanish for perfectly correlated *ν_α_*(*t*) and *ν_β_*(*t*). The product 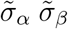 sets the slope of the linear decrease of 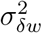 with *ρ*.

### The decay time of the autocorrelation function of width fluctuations is set by the mean volume growth rate and is only slightly perturbed by multiplicative noise (Fig. 4)

**Figure 4:**
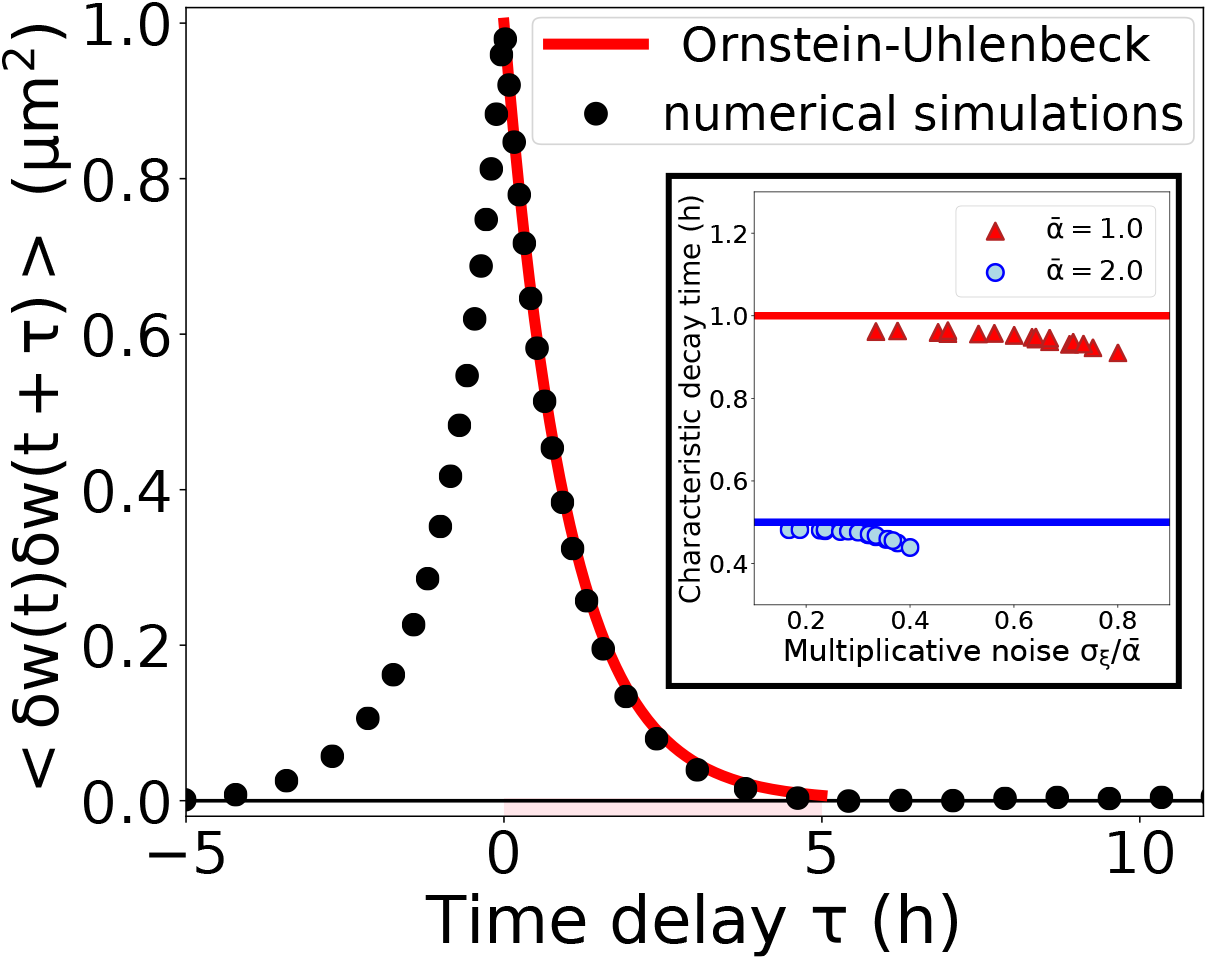
The decay time of the autocorrelation function of width fluctuations is set by the mean growth rate 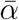 and is only slightly perturbed by multiplicative noise. Autocorrelation function of width fluctuations: the plot compares the average simulations (black circles) to the Ornstein-Uhlenbeck prediction (red line). Parameters: *α* = 1.0, *σ_ν_α__* = 0.05 (CV_*α*_ = 5%), *β* = 5.0, *σ_ν_β__* = 0.25 (CV_*β*_ = 5%). Inset: The exponential decay time is only mildly affected by the magnitude of multiplicative noise, measured in terms of its noise amplitude normalized by the average growth rate 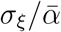. Parameters: 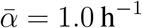, 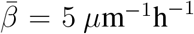 (red triangles and, for the Ornstein-Uhlenbeck prediction, line), 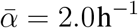, 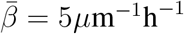 (blue circles and, for the Ornstein-Uhlenbeck prediction, line). The amplitude of the multiplicative noise was changed by simulating different combinations of *σ_ν_α__* and *σ_νβ_*.

Under the Ornstein-Uhlenbeck approximation (i.e. 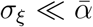) it is possible to obtain the autocorrelation function of width fluctuations at steady state analytically,

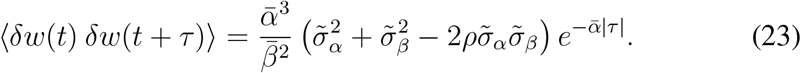

The characteristic time scale associated to width fluctuations is 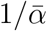, meaning that the average volume growth rate sets the time scale associated to how the system relaxes after width fluctuations or perturbations. This relaxation time scale is the same as that of mean width, and of the mean surface-to-volume ratio in a transition between environments (i.e. when 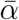 or 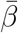) are changed [11]: in other words, the model predicts that width and surface-to-volume ratio are equilibrium variables that follow fluctuation-dissipation behavior.

The analytical result of Eq.23 is in a very good agreement with the computation of the autocorrelation from the numerical simulation of the complete system (Fig. 4). Only small deviations from the Ornstein-Uhlenbeck regime emerge by increasing the multiplicative noise (Fig. 4, inset).

### The time derivative of logarithmic length is a stationary variable whose fluctuations are driven by width fluctuations (Fig. 5)

**Figure 5:**
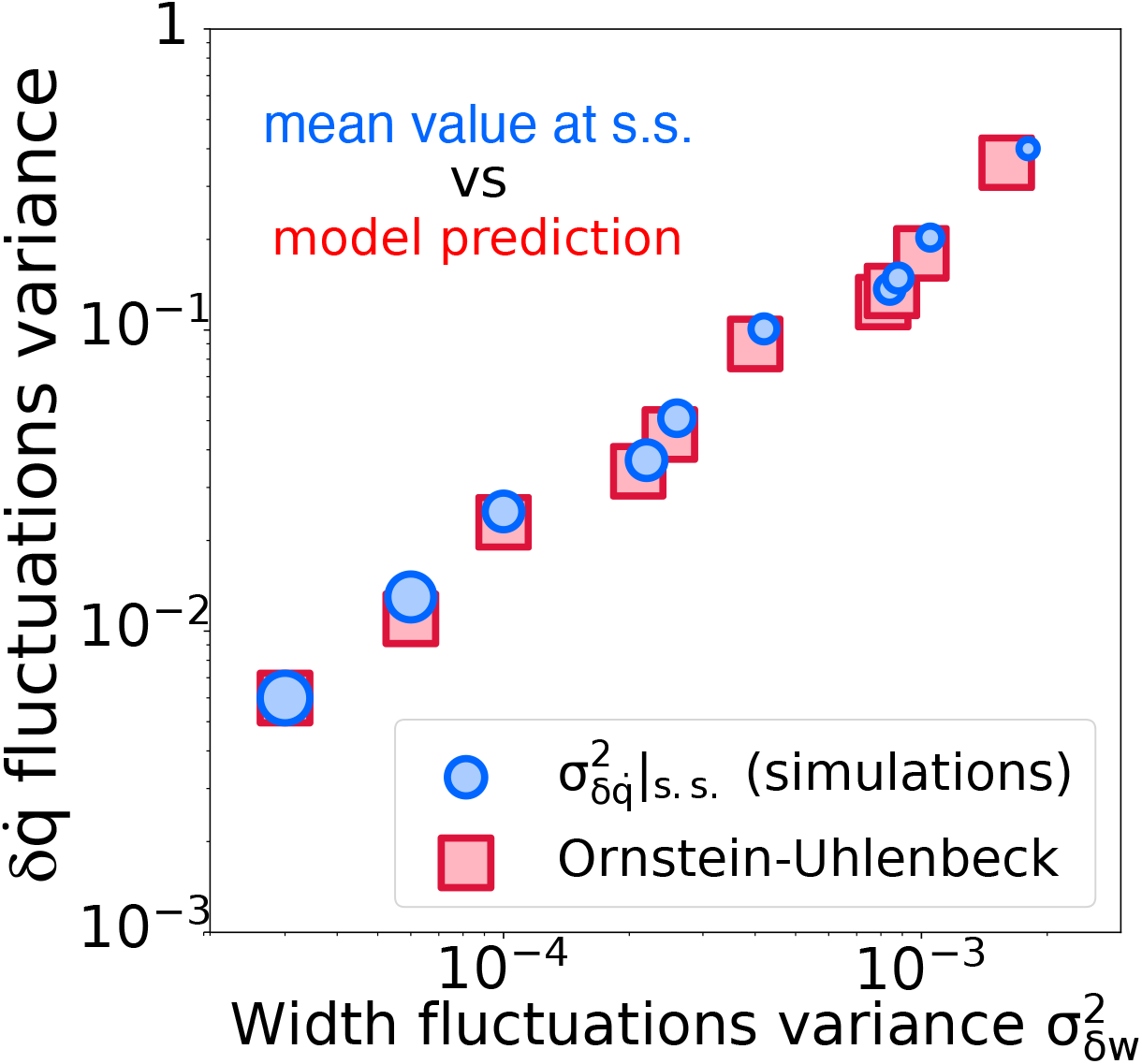
Fluctuations in the time derivative of logarithmic length at steady state are proportional to fluctuations in width under the Ornstein-Uhlenbeck regime. Comparison between the results of simulations (blue dots) and the analytical prediction under the Ornstein-Uhlenbeck regime (red squares). Different values of the width fluctuations variance are obtained by changing the amplitude of the noise on the growth rates *σ_ν_α__, σ_ν_β__*, while keeping constant their average value and their instantaneous correlation coefficient (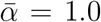, 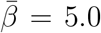, *ρ* = 0). The size of the blue dots is in inverse proportion to 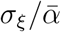, that is to the goodness of the Ornstein-Uhlenbeck approximation given the choice of the parameters.

While logarithmic length *q*(*t*) = log(*L*(*t*)/*L*_0_) is a non-stationary variable, its time derivative 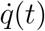 fluctuates around an average value 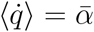. Hence, from Eq. 20 we can compute the analytical expectation for the variance of such fluctuations, at steady state and in the Ornstein-Uhlenbeck regime 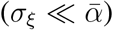:

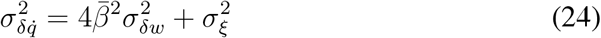

where 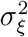 is provided by Equation 16 (please note that the additive noise in Eq. 20 corresponds to the multiplicative noise *ξ*(*t*) in Eq. 8). The numerical simulations are in agreement with the analytical prediction (Fig. 5).

### The shape of the width-log(length) cross-correlation function depends on the sign of the coupling between the volume- and the surface-specific growth rates

The cross-correlation functions between pairs of geometrical observables reveal their temporal hierarchy [16]. In our model, volume- or surface-specific growth rates *α, β* are delta-correlated by definition (Eq. 6c), while the cross-correlation functions between the rates and geometric observables such as width or logarithmic length are zero for negative delays by definition, as the rates are an input of the model, unaffected by other variables (Fig. 1).

In the Ornstein-Uhlenbeck approximation for the width fluctuations, specifically when

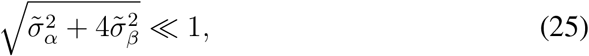

we can write the solution of the Langevin equation for the width fluctuations dynamics at steady state as

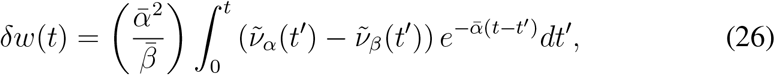

with *dt′* indicating the Itô integral. In such regime (*ξ* ≃ 0), the analytical expression of the cross-correlation functions between rates and geometric observables read:

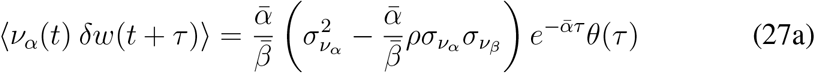

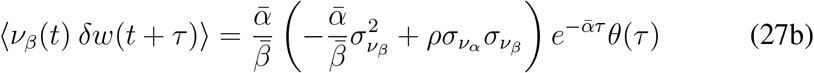

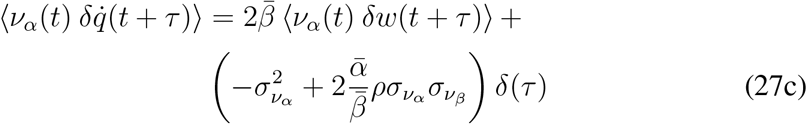

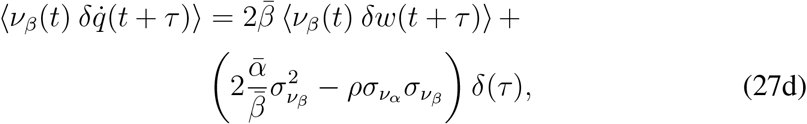

where *θ* and *δ* represent the Heaviside step and the Dirac delta functions, respectively. The time scale of the exponential decay of all the cross-correlation functions always equals 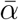.

The cross-correlations involving growth-rate fluctuations are not expected to be directly accessible in experimental data. However, the cross-correlations between fluctuations in cell width and those in length can in principle be measured. Since fluctuations in the time derivative of logarithmic length (Eq. 20) depend on an additive noise term (depending on both surface and volume synthesis), the steady-state cross correlation 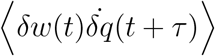 in the Ornstein-Uhlenbeck approximation for *δw* (Eq. 25) can be written as:

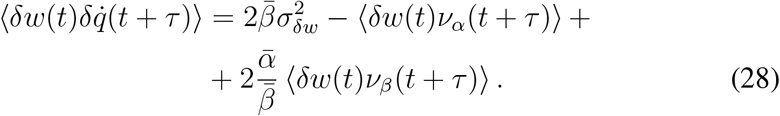

The numerical simulation and the Ornstein-Uhlenbeck prediction of 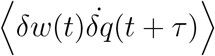 for different values of their coupling coefficient *ρ* are shown in Figure 6. We note that the coupling *ρ* between surface and volume synthesis can affect both quantitatively and qualitatively the time hierarchy between measured variables. Therefore, cross-correlations from data can in principle be used to infer the couplings and hence to make considerations about the relationship between the physiology of bulk vs surface accretion in single cells (Fig. 7).

**Figure 6:**
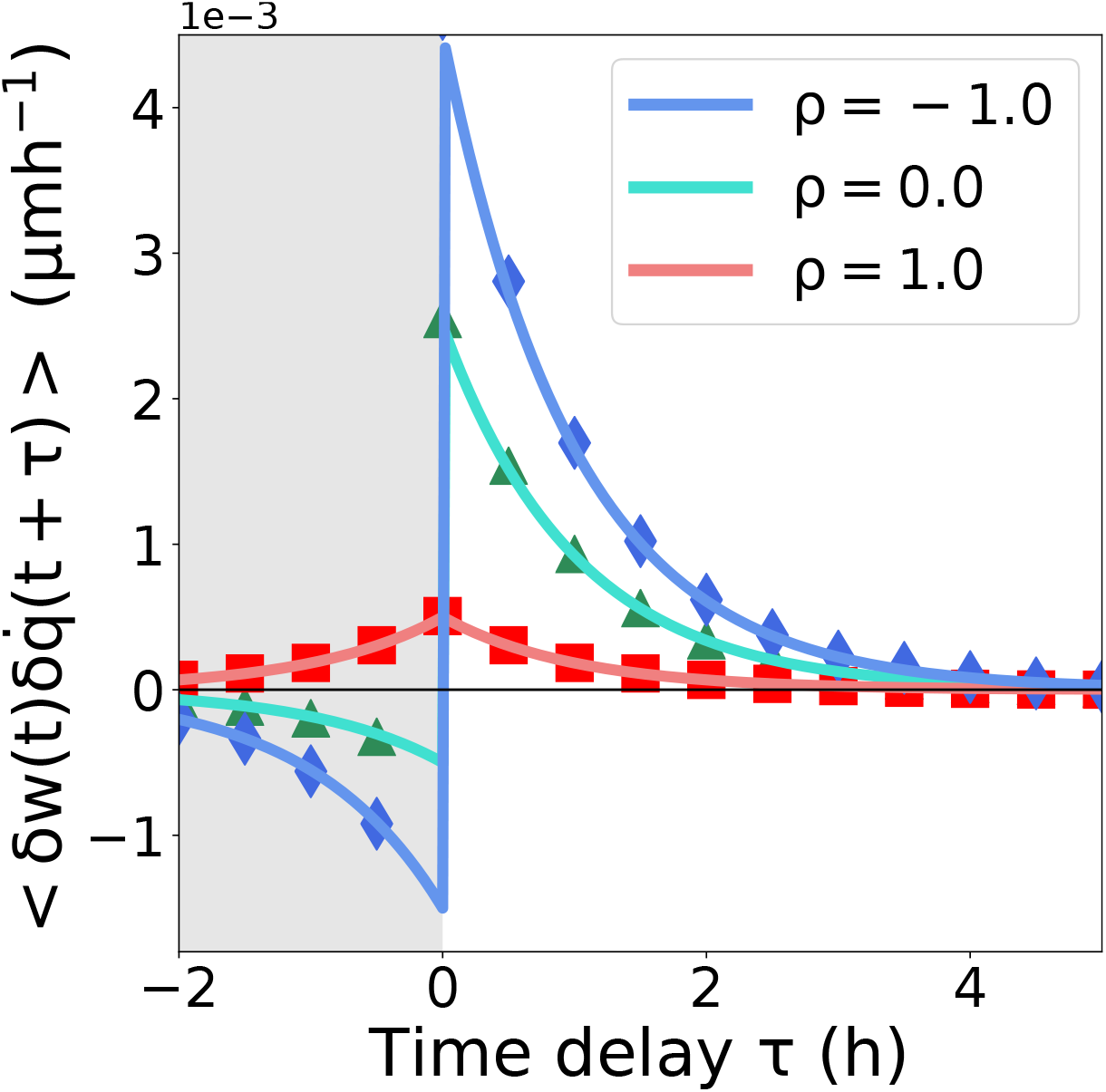
The coupling between volume- and surface-specific growth rates affects the cross-correlation function of fluctuations in width and in the time the derivative of logarithmic length. Unlike the decay exponent of the cross-correlation 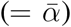, the value of 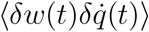 at zero delay is strongly affected by the correlation coefficient *ρ* between *α, β* fluctuations. With respect to the *ρ* ≤ 0 case, for certain combinations of the values of the coefficients of variation of *α, β*, increasingly positive values of the correlation coefficient *ρ* can result in a change of the sign of the correlation for negative delays. Parameters: *α* = 1.0, *σ_ν_α__* = 0.1, *β* = 5.0, *σ_ν_β__* = 0.25.

**Figure 7:**
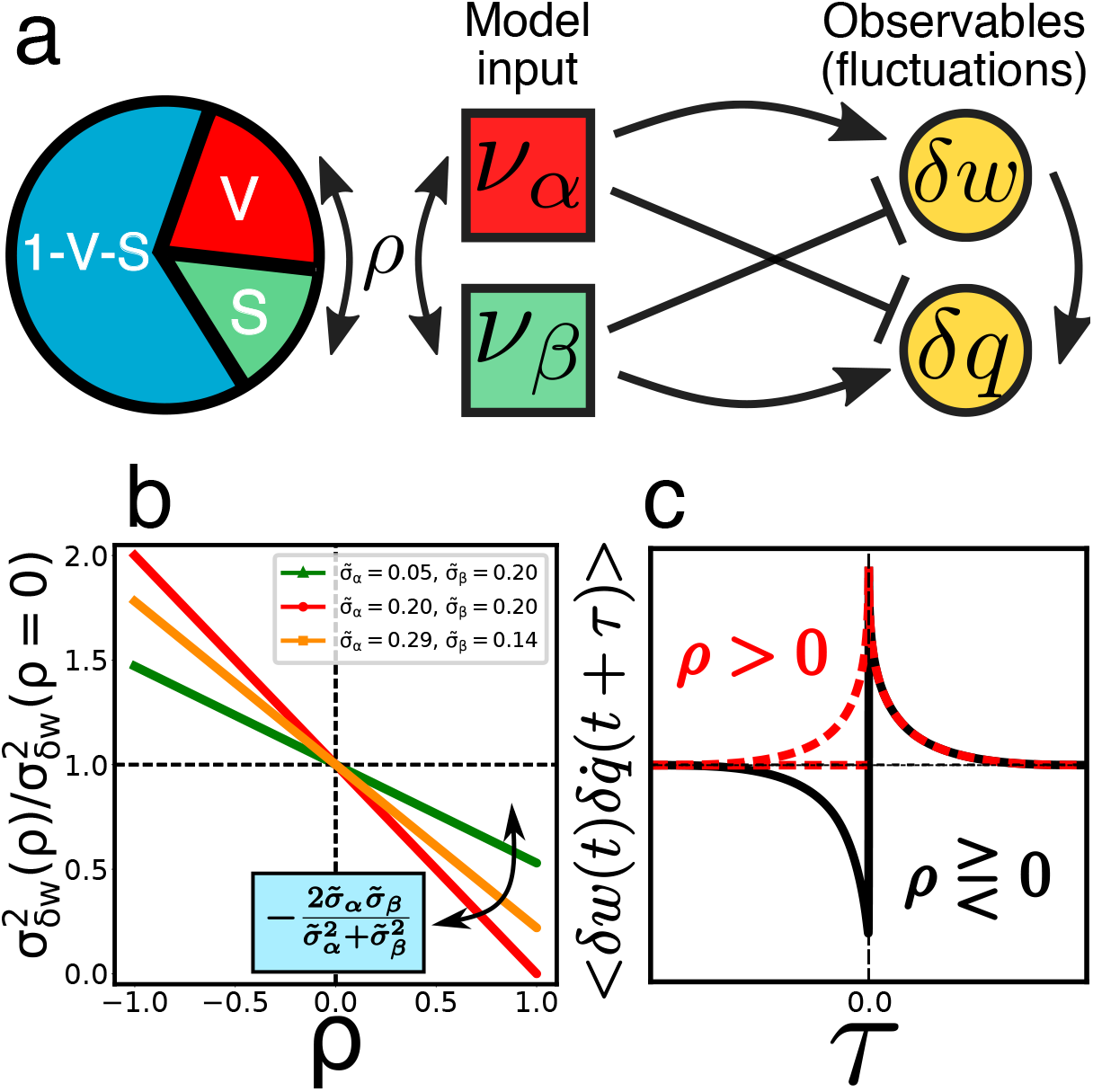
Cross-correlations in geometric features link to resource allocation of the cell. **(a)**: In the model, the parameter *ρ*, coupling surface and volume growth fluctuations, can be seen as a measure of the cooperativity (*ρ* > 0) vs antagonism/competition (*ρ* < 0) in the repartition of the resources between two specific sectors, such as those related to bulk (“V”) and surface (“S”) production. **(b)**: As previously seen, increasing *ρ* results in a decrease in cell-width fluctuations. With respect to the *ρ* = 0 case, the slope of the linear trend (well fitted by the prediction of the Ornstein-Uhlenbeck approximation) is a function of the coefficients of variation of *α* and *β* growth rates only. **(c)**: The shape of the width-log length dot cross-correlation function for *τ* < 0 can be used to isolate scenarios of positive vs zero- or negative correlation between surface and volume “sectors”. In particular, for negative delays it would be impossible to observe positive or zero correlation between width fluctuations *δw* and fluctuations in the derivative of log length 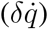 in the case the “S” and “V” sectors competed for the same limiting resources (*ρ* < 0).

## 4 Discussion

While it is evident that both volume and surface accretion must affect bacterial shape homeostasis, the role of the coupling between the two remains an open question. From a cellular metabolism perspective, to a first approximation volume growth relates to the production of all proteins and metabolites increasing cellular biomass, while surface growth relates to those proteins specifically needed for building the cell wall and the outer membrane [19]. Since both the energy and the ribosomes required for both processes are usually limiting factors [15], the bacterial cell has to perpetually solve a resource allocation problem to maintain the right balance between bulk and surface-specific synthesis and thus maintain shape homeostasis between successive divisions.

We developed a single-cell phenomenological model, which is analytically solvable and provides experimentally testable predictions. For an average cell, the model is identical to the Harris and Theriot model, validated in both gram positive and negative bacteria such as *E. coli, C. crescentus* and *L. monocytogenes* [11] despite of differences in the cell-wall and membrane composition [20, 21]. An important assumption of the model is considering volume- and surface-related growth rates as inputs of the model (their fluctuations are *a priori*, without a feedback with other dynamic variables). A further, important assumption is that of constant density: this is needed if we want to identify volume synthesis with bulk biosynthesis for single cells. Finally, we chose not to investigate the role of cell cycle and division, so that our predictions should be in principle relatable to what happens to growing bacteria between two successive divisions.

Crucially, the model contains a correlation between surface and volume growth-rate fluctuations, quantified by the single parameter *ρ*, describing the coupling between different biosynthetic sectors. We find that this parameter affects only some of the measurable observables. In particular, the model predicts a linear, negative dependency between *ρ* and the variance of width fluctuations (Fig. 3). The cross-correlations functions involving rates’ fluctuations (e.g., Eq. 27a) and between pairs of shape-related observables (Fig. 6) are also qualitatively affected by the value of *ρ*. On the contrary, the coupling *ρ* does not affect the decay times of the auto- and cross-correlation functions, set by the average growth rate 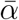 only (see Eqs. 23 and 27a-d, and Figs. 4 and 6). We interpret (Fig. 7a) a positive *ρ* as cooperativity between the processes related to the synthesis of all vs surface-specific “building blocks”. Conversely, negative values of *ρ* would correspond to competition for the limiting resources between the surface and bulk synthesis (for example as a result of the exposure to lack of nutrients or-stress).

Our model suggests the possibility to infer the sign of *ρ* from measurable cross-correlations (for which we reported the analytical expressions), which would be of interest to effectively investigate the physiology behind the problem of shape homeostasis in bacteria. Indeed, cross-correlations provide with a hint of the temporal hierarchy between couples of fluctuations time series, thus suggesting possible causality links between them [16]. We expect the rates-observables cross-correlation functions (Eqs. 27a-d) to be of problematic experimental investigation, unlike the width-width autocorrelation function (Eq. 23), which should be a standard measurable quantity. However, the effect of different *ρ* on this autocorrelation function is only quantitative, meaning that the determination of the sign (and value) of *ρ* would require measuring the growth rates and their coefficients of variation as well.

A safer route to extract information on the sign of *ρ* would then be to rely upon the qualitative change in the width-log length dot cross-correlation function (Eq. 28), which we expect to be measurable, too. In the Ornstein-Uhlenbeck regime and at steady state, the cross-correlation between fluctuations in width and those in the time derivative of logarithmic length (which are both stationary processes) is given by the sum of the variance of the width fluctuations and a linear combination of the cross-correlation between rates’ and width fluctuations. Crucially, the model predicts a positive or zero cross-correlation for *τ* < 0 (that is, a positive correlation between fluctuations in logarithmic length and width fluctuations later in time) only for positive values of *ρ* (Fig. 7c).

We believe that this route to extract physiological information from shape fluctuations is feasible in general, for example with models allowing for a higher number of measured variables. In particular, recent studies tackled the problem of density homeostasis through the mathematical modelling of population data [22]: our stochastic model provides one possible mathematical framework for the study of density fluctuations.

